# Unique features of the gut microbiome characterized in animal models of Angelman Syndrome

**DOI:** 10.1101/2022.07.05.498914

**Authors:** Ulrika Beitnere, Brayan Vilanova-Cuevas, Sarah G Christian, Clint Taylor, Elizabeth L Berg, Nycole A Copping, Scott V. Dindot, Jill L Silverman, Mélanie G Gareau, David J Segal

## Abstract

A large subset of patients with Angelman syndrome (AS) suffer from concurrent gastrointestinal (GI) issues, including constipation, poor feeding, and reflux. AS is caused by the loss of ubiquitin ligase E3A (*UBE3A*) gene expression in the brain. Clinical features of AS, which include developmental delays, intellectual disability, microcephaly, and seizures, are primarily due to the deficient expression or function of the maternally inherited *UBE3A allele*. The association between neurodevelopmental delay and GI disorders is part of the increasing evidence suggesting a link between the brain and the gut microbiome via the microbiota-gut-brain (MGB) axis. To investigate the associations between colonization of the gut microbiota in AS, we characterized the fecal microbiome in three animal models of AS containing maternal deletions of *Ube3A*, including mouse, rat, and pig, using 16S ribosomal RNA amplicon sequencing. Overall changes in the microbial composition of all three animal models of AS in both the phylum and genus levels of bacterial abundance were identified. Specific bacterial groups were significantly increased across all animal models, including: *Lachnospiraceae Incertae sedis, Desulfovibrios sp*., and *Odoribacter*, which have been correlated with neuropsychiatric disorders. Taken together, these findings suggest that specific changes to the local environment in the gut are driven by a *Ube3a* maternal deletion, unaffected by varying housing conditions and are prominent and detectable across multiple small and large model species. These findings may begin to uncover the underlying mechanistic causes of GI disorders in AS patients and provide future therapeutic options for AS patients.

**IMPORTANCE:** Angelman syndrome (AS) associated gastrointestinal (GI) symptoms significantly impact quality of life in patients. Using AS models in mouse, rat, and pig, AS animals showed impaired colonization of the gut microbiota compared to wild type (healthy) control animals. Unique changes in AS microbiomes across all three animal models may be important in causing GI symptoms and may help to identify ways to treat these comorbidities in patients in the future.

## INTRODUCTION

Angelman syndrome (AS) is a rare (1 in 15,000 births) genetic neurodevelopmental syndrome caused by the loss of maternally inherited ubiquitin ligase E3A (*UBE3A*) gene expression in mature neurons of the brain (1, 2). The paternal copy of *UBE3A* is expressed in most peripheral organs, potentially leading to haploinsufficiency in these tissues. However, due to brain-specific imprinting, paternal *UBE3A* is silenced in the central nervous system (CNS) by a long non-coding antisense transcript (*UBE3A-ATS*), resulting in a complete loss of UBE3A expression in the brain (3). This genetic configuration in AS leads to microcephaly, severe developmental delays, deficiencies in expressive communication, typical facial appearance, deficits in movement and coordination, hypotonia, generalized epilepsy, sleep disturbances, and other characteristic behaviors, such as frequent smiling and laughter (4). In addition to the effects on neurodevelopment, many caregivers report gastrointestinal (GI) issues in AS patients. Particularly, children with AS are often reported as poor feeders due to hypotonia of the throat (5) and have a high rate of constipation (6). Despite this strong association of GI disorders in AS patients, the mechanisms underlying this remain largely unknown.

The microbiota-gut-brain (MGB) axis represents the bidirectional communication pathways that connect the gut-microbiota to the brain and modulate behavior (7). Studies using germ-free (GF) mice have identified multiple behavioral impairments, including cognitive deficits (8) and anxiolytic behaviors (9), compared to colonized controls, supporting a role for gut microbes in maintaining these behaviors. Regulation of behaviors by gut bacteria might occur via a combination of multiple pathways, including endocrine signaling through hormones and neuro-active metabolites, as well as signaling via the immune system and vagus nerve. Colonization of the gut microbiome begins at birth and plays a critical role in building a healthy gut, shaping immune processes and neurodevelopment (10, 11).

The disruption of microbial communities in the GI tract has been implicated in a number of different neurodevelopmental and neurodegenerative disorders, such as autism spectrum disorders (12, 13), Alzheimer’s disease (14, 15), Parkinson’s disease (16–18), depression (19), amyotrophic lateral sclerosis (20–22), schizophrenia (23, 24), and attention-deficit/hyperactivity disorder (25). Monogenetic neurodevelopmental disorders may also exhibit changes in microbial composition that may explain some of the GI symptoms seen in subsets of patients. For example, Rett Syndrome, a severe and progressive X-linked neurological disorder affecting mainly females due to mutations in the *MECP2* gene, has a strong association with GI dysfunction, including intestinal dysbiosis which is characterized as a disruption to the bacterial homeostasis (26–28). Relevant to the work herein, the same region on chromosome 15 that is affected in AS, causes Prader-Willi syndrome (PWS) when the paternal contribution of genes on chromosome 15 is lost. Although clinically distinct from AS, individuals with PWS exhibit obesity, hyperphagia, and reduced metabolic rate, in the context of an altered microbiome (29–31). The frequency and scope of GI illnesses in AS, however, have never been studied and the diagnostic consensus estimates that the prevalence may affect between 70% of individuals with AS (6). GI problems in AS were reviewed using medical records of 163 individuals with AS with different genetic subtypes and characterized, identifying at least one GI dysfunction in most patients (6). The two most common dysfunctions were constipation and gastroesophageal reflux disease (GERD). Other GI problems reported included cyclic vomiting episodes, difficulty swallowing, excessive swallowing, and eosinophilic esophagitis (6). Despite this prevalence of GI symptoms in AS patients, a 16S ribosomal RNA sequencing study in AS patients has not been performed yet.

To study the potential impacts of AS on the brain on the gut microbiota, three AS animal models that lack maternal UBE3A expression, similar to humans, were compared. The mouse model contains an inserted nonsense mutation in exon 2 od the mouse *Ube3a* gene (32), whereas the rat (33–35) and pig models have a full gene deletion from the use of CRISPR/Cas9 nucleases flanking the *Ube3a* gene. We compared the global gut microbial community (alpha and beta diversity), as well as the microbial composition at the taxonomic phylum and genus level in all three animal models. In addition, to better understand the underlying metabolic processes affected by changes in the microenvironment of the gut microbiota in genetic animal models of AS, an inference of metabolic pathways based on the bacterial microbiome was performed. While similar changes were seen at the phylum level in AS animals compared to controls, distinct patterns were observed in each species.

## METHODS

### Animals

Mice used in the study were B6.129S7-*Ube3a*^*tm1Alb*^/J (Jackson Laboratory strain #:016590) (32). Wild type (WT) C57B6/J littermates served as controls. The recently characterized *Ube3a*^m*/p+^AS rat model contained a full 90-kb deletion of the maternal *Ube3a* gene on Sprague-Dawley background (33, 34), with WT littermates serving as controls. The AS pig model (*Sus scrofa*) contained a full 97-kb deletion of the maternal *UBE3A* gene on a mixed Yorkshire/Landrace background (S.V.D., personal communication). Littermates of both genotypes were housed together for all species.

### Fecal sample collection

In total, 41 mice, 26 rats, and 8 pig fecal samples from both sexes were collected for this study (**Table 1**). Mice were 13-25 weeks of age, rats were 8 weeks of age, and the pigs were 17 weeks old, at the time of the fecal sample collection. All animals were housed in appropriate light/dark conditions and fed standard food/water according to the model’s dietary needs. Mice and rats received Teklad global 18% protein rodent diets 2918 (Envigo, Hayward, CA, USA). Pigs received MG Pig Starter 20% - Gen 2.0 for pigs weighing up to 44 pounds and for adults: MG Hog Pellets (M-G, INC. Feed Division, TX, USA).

**Table #1.**
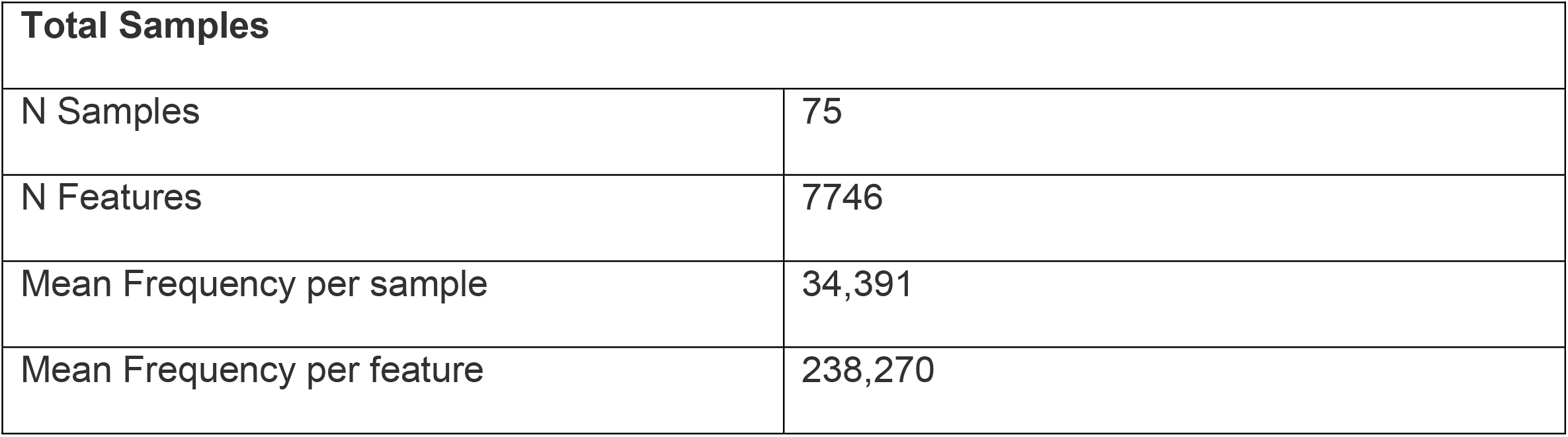
Sample and Feature Sequence Frequency.

| Total Samples |  |
| --- | --- |
| N Samples | 75 |
| N Features | 7746 |
| Mean Frequency per sample | 34,391 |
| Mean Frequency per feature | 238,270 |

Fecal samples from mice and rats were collected at the University of California, Davis. To collect the fecal samples, animals were placed individually in an empty sterile cage for 5 min and freshly dropped fecal pellets were collected aseptically into sterile tubes using sterile pipette tips or sterile tweezers. Samples from pigs were collected at Texas A&M University. The samples were collected after euthanasia using a sterile fecal loop into sterile tubes and stored at −80 C until further processing.

### 16S Illumina sequencing

DNA was extracted from 20 - 40 mg fecal matter using the QIAamp Powerfecal Kit (Qiagen). The library preparation and sequencing was performed by the Host Microbe Systems Biology Core at UC Davis using primer pair 341F and 806R in 300 bp paired-end run for the V3-V4 hypervariable regions of the 16S rRNA on an Illumina MiSeq (Illumina, San Diego). The 16S rRNA Raw FASTQ sequence files were deposited and processed in QIITA (36) using per-sample FASTQs with a Phred offset of 33, min_per_read_length_fraction of 0.75 and default parameters for error detection using Split libraries FASTQ. Sequences were trimmed to 250 bp and possible errors of sequencing were filtered using DEBLUR with default parameters. Reference operational taxonomic units (OTUs) were defined using the SILVA reference database with a minimum similarity threshold of 97% and corresponding taxonomy assignment using the default parameters in QIITA. Singletons (OTUs with less than three reads), sequences matching chloroplasts, mitochondria, and unassigned sequences were removed from downstream analyses followed by a rarefaction to the minimum library size, which was 21907. The main variable utilized for analysis was genotype, WT control or AS, with each species assessed individually or compared to each other in a single group analysis.

Alpha diversity, beta diversity, and taxonomic composition plots were built using R’s ggplot2 package (37, 38). Beta diversity analyses of microbial communities were performed by computing the pairwise Bray–Curtis distances (39) between samples and plotted using non-metric multidimensional scaling (NMDS). To determine the significance of the dispersion between the samples, the results of analysis dissimilarities were calculated directly from the distance matrix with an ADONIS. Alpha richness (Chao1: estimated number of OTUs), and diversity (Shannon: index of equitability) (40) indexes were used to establish significant differences between the genotypes and animal models, which were assessed with the non-parametric Wilcox test. Significance was defined as p<0.05. To establish significant differences between specific OTUs, we applied a fold change analysis using the Deseq2 pipeline (41) to visualize possible bacterial biomarkers related to the genotype in all the samples, in the separate models.

### PICRUST metabolic analysis

Metabolic pathway inference analyses were performed using PICRUST2 software (42). A pathway level-inference analysis was performed, where MetaCY pathways are inferred using enzyme classification number of abundances. Output used for analysis is composed of an unstratified (sum of all sequences contributing by OTUs) pathway abundance table. Analysis was performed using Metaboanalyst software (43) where pathways were filtered by the mean intensity and log transformation of the abundance counts. Subsequently, the ward clustering method was used and Euclidean distances to group heatmaps presented only significantly different pathways identified. Significance was calculated using a t-test or ANOVA as appropriate.

## RESULTS

### The global microbial community structure in AS animal models is specific to each species

Richness and diversity measures across all species were performed. These findings indicate that the mouse model is much less rich in total bacterial numbers in comparison to both pig and rat models (**Fig 1A**). In contrast, the diversity index shows that both mouse and pig models are similar in diversity, whereas the rat model is significantly higher in diversity (**Fig 1A**). Significant dispersion based on animal model (***p < 0.001) was observed, however there is no significant separation based on genotype (p = 0.7; **Fig 1B**).

**Figure 1.**
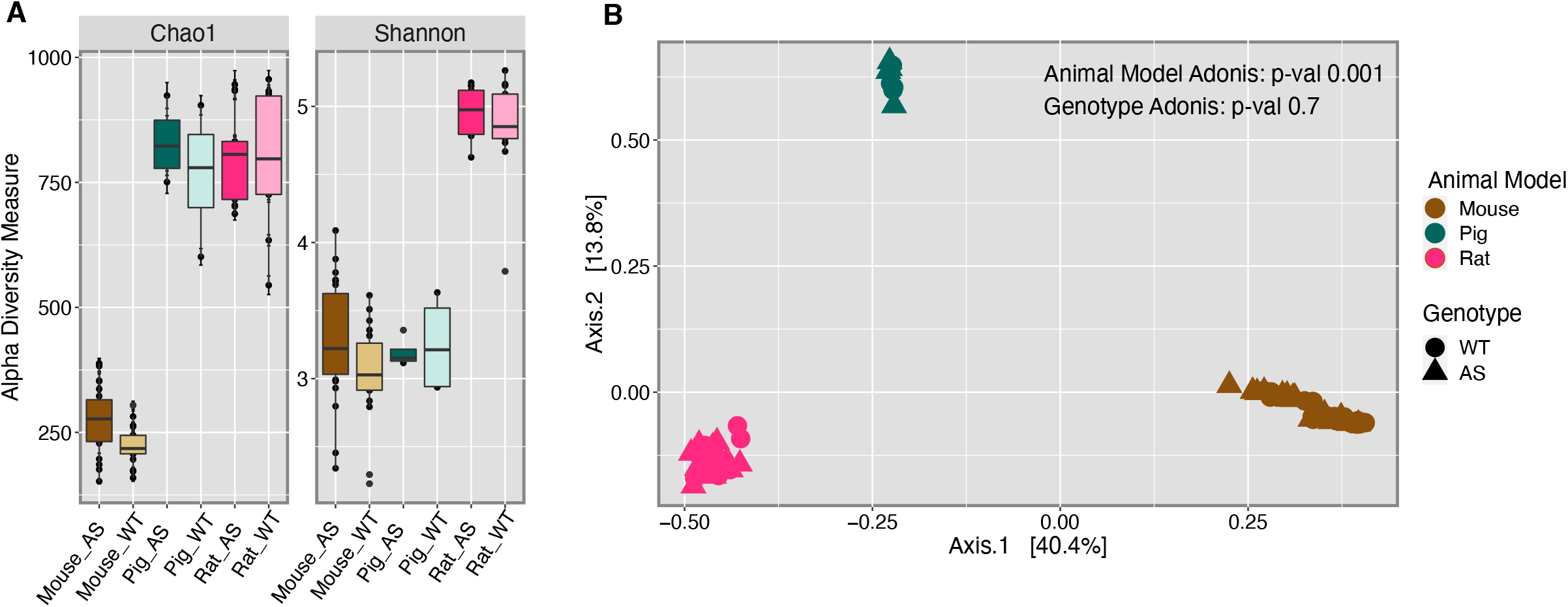
Microbial community structures differ between the animal models. Alpha diversity analysis (**A**) presents both richness (Chao1) and diversity (Shannon) indexes, and Kruskal-Wallis pairwise comparison was performed to determine the difference between the overall animal models. Rats are significantly richer (*p* = 6.992939e-12) and diverse (*p* =l 8.370003e-12) than mice. Rats are also more diverse than pig (*p* = 2.417261e-05), while pigs are richer than mouse (*p* = 9.155458e-06). Beta diversity analysis using Bray-Curtis dissimilarities index shows significant dispersion of the samples by animal model (Adonis *p* = 0.001).

Individual animal model analysis was performed to assess specific differences in the microbial community structure between WT vs. AS for each animal species. The microbial community structure of individual animals relative to the genotype identified no significant differences between genotypes across all three species (**Fig 2A,C, E**). Alpha diversity analysis exploring the richness and diversity index of the individual species identified significant differences in mice, with both Chao1 and Shannon indexes supporting a richer (p < 0.01) and more diverse (p < 0.05) microbiome in AS mice compared to WT controls (**Fig 2B**). In contrast, no significant differences were identified in either pig or rat between AS and WT animals (**Fig 2D,F**).

**Figure 2.**
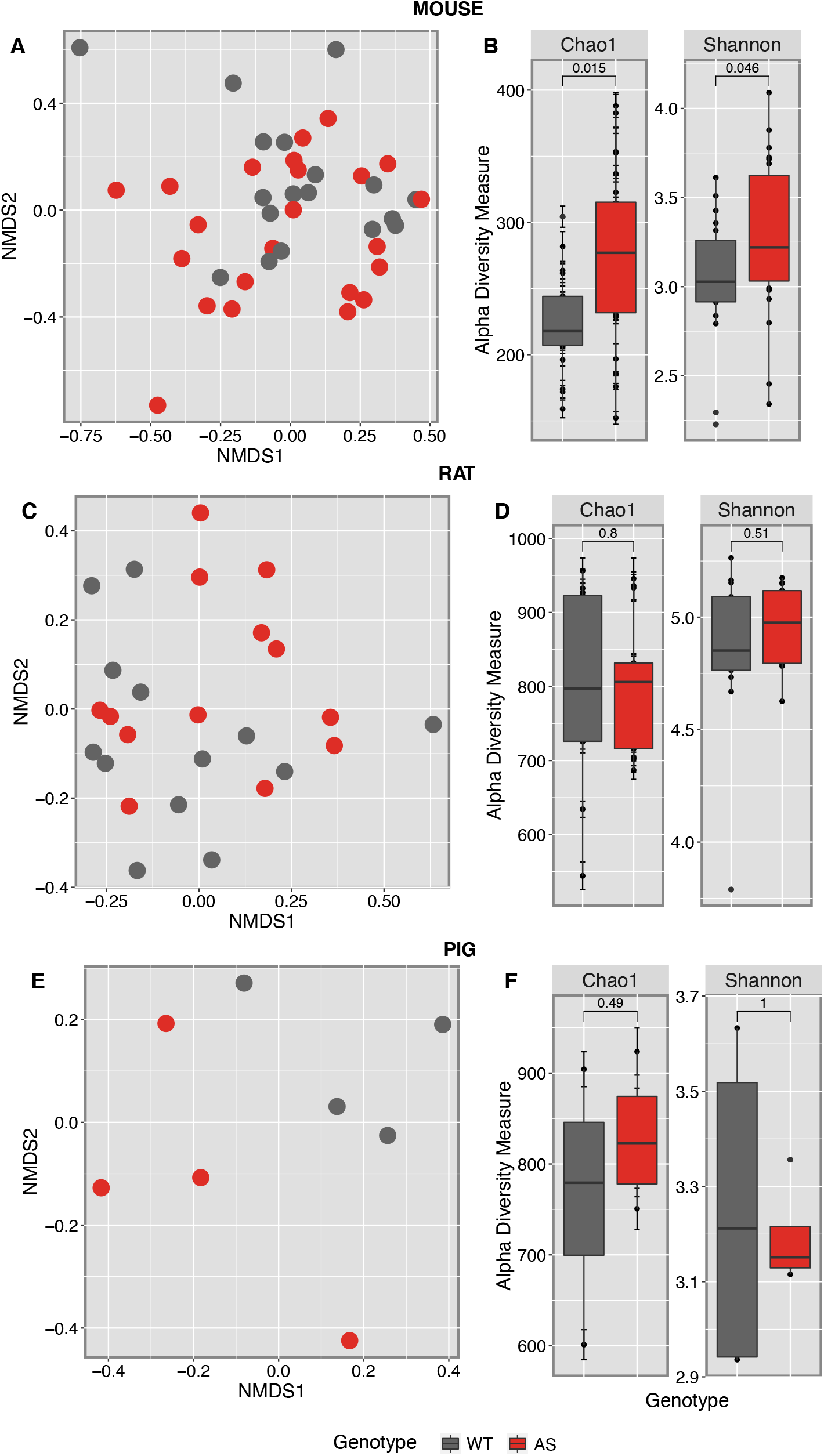
Individual animal models’ overall gut microbial composition structure is not altered by AS genotype. Bray-Curtis dissimilarities index was used to analyze the dispersion of samples using an NMDS visualization. The mouse model presents no significant dispersion of the samples (**A**), but alpha diversity shows AS mice are richer (*p* = 0.015) and more diverse (*p* = 0.046) than WT (**B**). Rat and Pig models show no significant dispersion (**C & E**), and no difference in alpha richness or diversity indexes (**D & F**).

### Overall microbial composition differences in multiple AS animal models

To characterize the differences between the most abundant operational taxonomic units (OTUs) in each genotype, the relative abundance of all OTUs was calculated and only phyla and genera with a minimum of 1% relative abundance were included in the analysis. Phylum level differences in relative abundance are consistent between the AS and WT in all animal models (**Fig 3**). Across the three AS animals, a reduction of *Firmicutes* (green) and an increase of *Bacteroidota* (red) were observed compared to WT controls, representing the major phyla for all three animal models (**Fig 3A-C**). In contrast, an increase of *Actinobacteriota* was identified in both AS mice and pigs, which was reduced in the AS rats in comparison to WT (**Fig 3B**). These differences at the phylum level suggest that the AS genotype disrupts the abundance of the most highly abundant phyla in the gut, with similar changes observed across multiple species.

**Figure 3.**
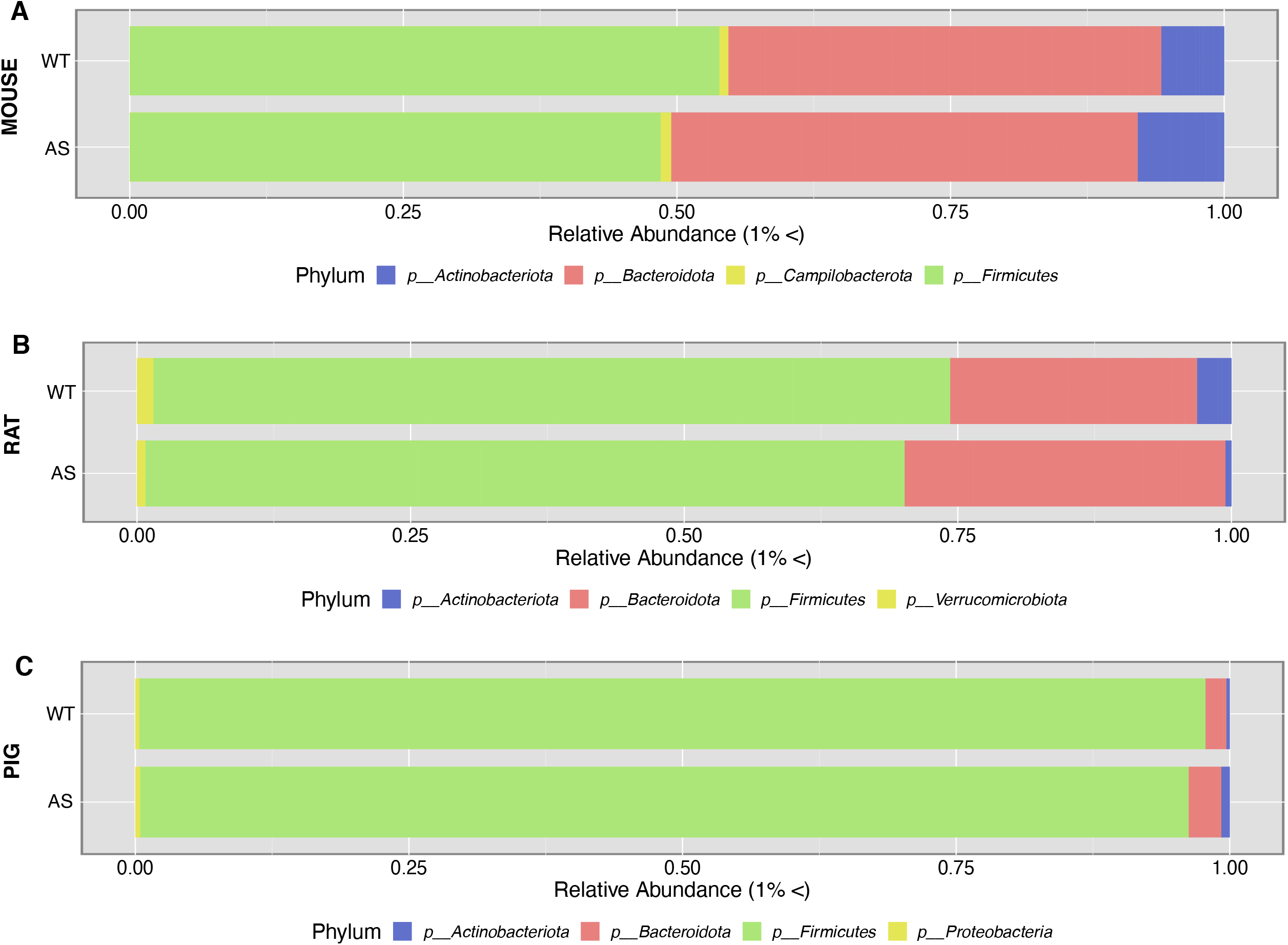
Phylum level differences between AS and WT by animal model. All panels represent phyla composition present above 1% relative abundance. A comparison of overall phyla composition between the animal models (mouse [**A**], rat [**B**], pig [**C**]) and genotype (AS and WT) presents a similar decrease of Firmicutes and an increase of Bacteroides across all species.

At the genus level, all AS animals demonstrated differences in abundance when compared to their respective WT control. For example, in AS mice, an increase of *Bacteroides, Coriobactericeae UCG-002, Faecalibaculum, Helicobacter, Incertae Sedis, Lachnospiraceae NK4A136 group and UCG-006, Marvinbryantia*, and *Turicibacter* was observed in comparison to WT control mice (**Fig 4A**). This was accompanied by a reduction of *Lactobacillus*, and *Dubosiella* (**Fig 4A**). In AS rats, an increase in *Bacteroides, Blautia, Gastranaerophilales, Monoglobus, Nocardia, Roseburia*, and *Ruminococcus* was seen (**Fig 4B**). This was coupled with a decrease in *Akkermansia, Eubacterium ventriosum group, Tepidibacter*, and *Lachnospiraceae UCG 001* compared to WT controls (**Fig 4B**). Lastly, AS pigs had increased *Subdoligranulum, Tepidibacter, Treponema, Faecalibacterium, Blautia*, and *Butyricicoccus*, whereas there was a decrease in *UCG-005, Costridium sensu stricto1, Fibrobacter, Monoglobus*, and *Streptococcus* compared to WT control pigs. While species-level characterization is preferable to establish the specific role of each organism in the gut, the genus characterizations can provide an important picture of how the genetic impairment affects the microbial composition (44). Taken together, the analysis of all the animal models shows differences between the AS and WT animals at the genus level, but these differences do not overlap across models, in contrast to the findings at the phylum level, highlighting the difference in composition across each species.

**Figure 4.**
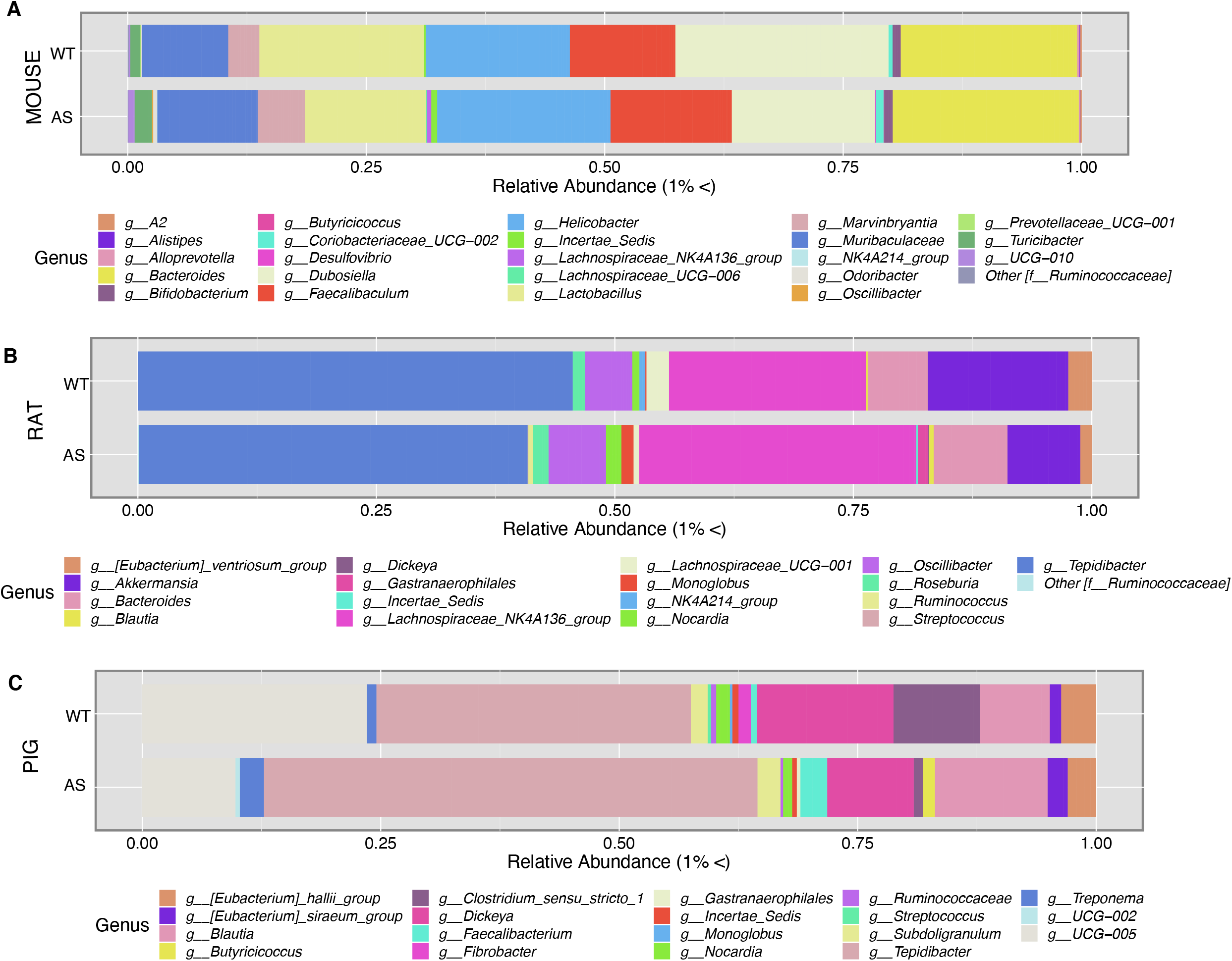
Genus level differences between AS and WT by animal model. Relative abundance of genera above 1% shows different genus compositions across all animal models. Mice **(A)** show a decrease in *Dubosiella* and *Lactobacillus* and an increase in *Helicobacter and Bacteroides* in AS vs WT. In rats **(B)**, we see an increase in the *Lachnospiraceae NK4A136 group* and a decrease in *Akkermansias* in AS vs WT. Finally, in the pig model **(C)**, we see an increase in *Tepidibacter* and *Blautia*, a decrease in *Clostridium sensu stricto 1*, and *Lachnospiraceae UCG-005* in AS vs WT.

### Bacterial biomarkers identified across and within AS models

Due to the wide differences in genus-level taxonomic composition found between the different animal models, a fold change (Deseq2) analysis was performed by genotype and separated by animal model. This analysis helps establish specific high and low abundant taxon that are differentially abundant within the microbial ecosystem according to genotype. Analysis of AS vs WT across all three animal models identified a differential abundance of *Desulfobacterota, Bacteroidota* and *Firmicutes* genera based on genotype. Genus level differences showed a higher prevalence of *UCG-010, Incertae Sedis, Desulfovibrio, Odoribacter*, and *Butyricicocaceae* family members in AS animals (**Fig 5A**). In contrast, WT animals had a differential abundance of *Clostridium sensu stricto, NK4214 group*, and *Lachnospiraceae UCG 001* (**Fig 5A**).

**Figure 5.**
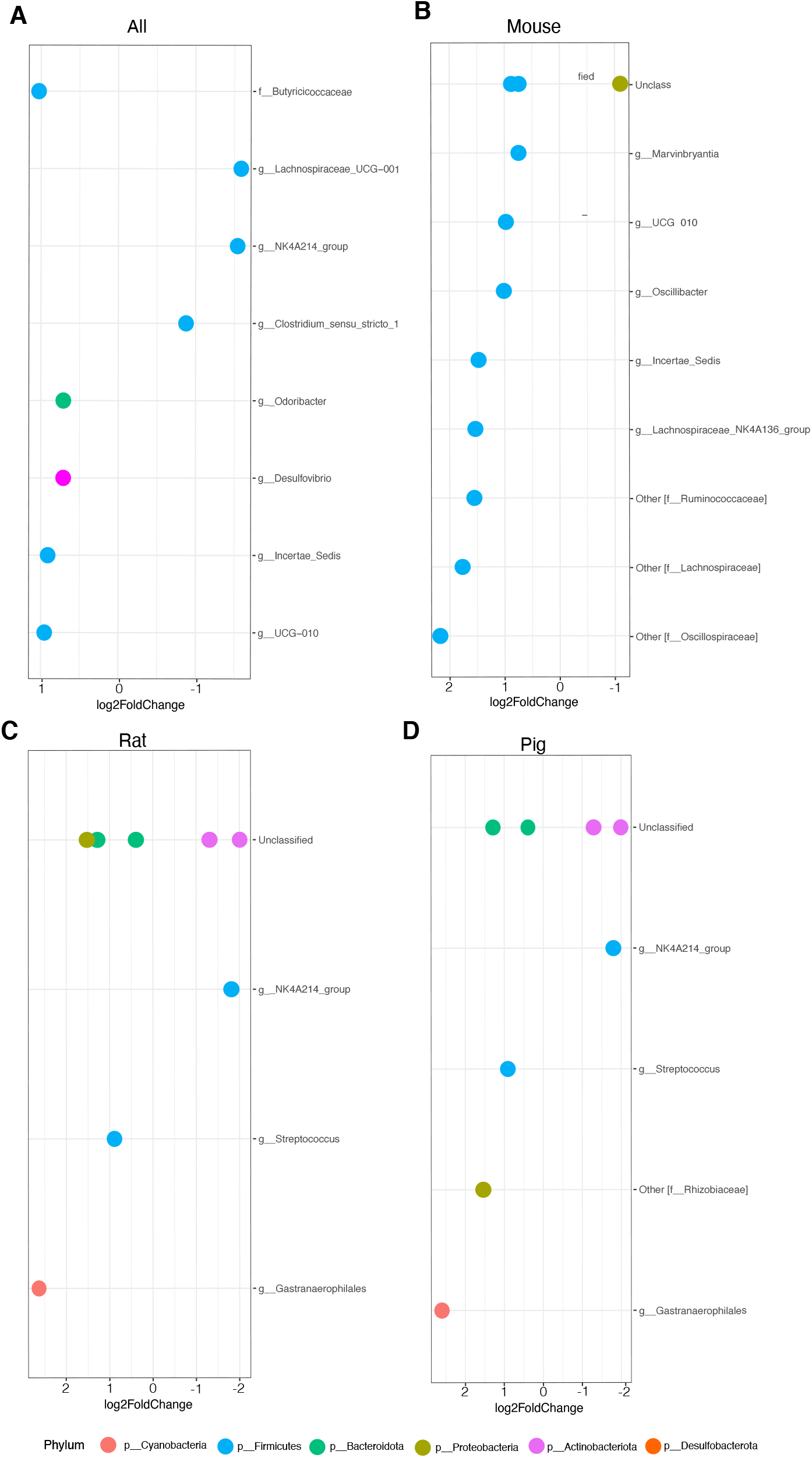
Fold change analysis shows significantly different bacterial groups associated with AS. This analysis only considered bacteria that showed significant differences between the genotype (*p* < 0.05). (**A**) Overall differences between genotypes across the microbiome of all animal models. Values above 0 are correlated with AS and values below are correlated with WT. (**B**,**C**,**D**) The bacterial groups associated with AS in individual animal models (mouse, rat, and pig).

While these differences account for the genotype in all the animal models, there are also innate differences between the gut microbiome of all three species. Therefore, fold change analysis was performed in all the models individually by genotype. In mice, a differential abundance of *Incertae Sedis, Oscillibacter, UCG 010, Marvinbryantia, Lachnospiraceae NK4A136 group, and other unclassified genera pertaining to Oscillospiraceae, Laachnospiraceae*, and *Ruminococaceae* families were identified in the AS model compared to WT controls (**Fig 5B**). In the rat and pig models, fewer bacterial genera were differentially abundant in AS animals, these being *Gastranaerophilales, Streptococcus, and Rhizobiaceae* family (only in pig), and some unclassified *Bacteroidota* and *Proteobacteria* (only in rat) phylum-level bacterium compared to WT controls (**Fig 5C,D**). WT rats and pigs presented with *NK4A214* group and unclassified *Actinobacteria* as being highly prevalent in compared to AS animals (**Fig 5C, D**). These findings suggest similarities between pig and rat microbial ecosystems and illustrate how the AS genotype affects the intestinal bacterial community.

### Differential bacterial metabolic pathways identified in each animal model of AS

To better understand the underlying metabolic processes affected by changes in the microenvironment of the gut microbiota in genetic animal models of AS, an inference of metabolic pathways based on the bacterial microbiome was performed using PICRUST. The metabolic pathway analysis was performed for both individual species and the animal models combined, which allowed visualization of important pathways that play a role in each animal model and pathways that are impacted specifically due to genotype. The combined model analysis shows that the microbiota in AS animals have higher activity in processes such as glycolysis, lactic fermentation, glycan building blocks, nucleoside biosynthesis, vitamin B1 synthesis, vitamin B5, and CoA biosynthesis, and urate production and accumulation (**Fig 6A**) compared to WT controls. These results suggest changes in the metabolic pathway activity based on genotype, yet preexisting differences based on each animal model can introduce variability.

**Figure 6.**
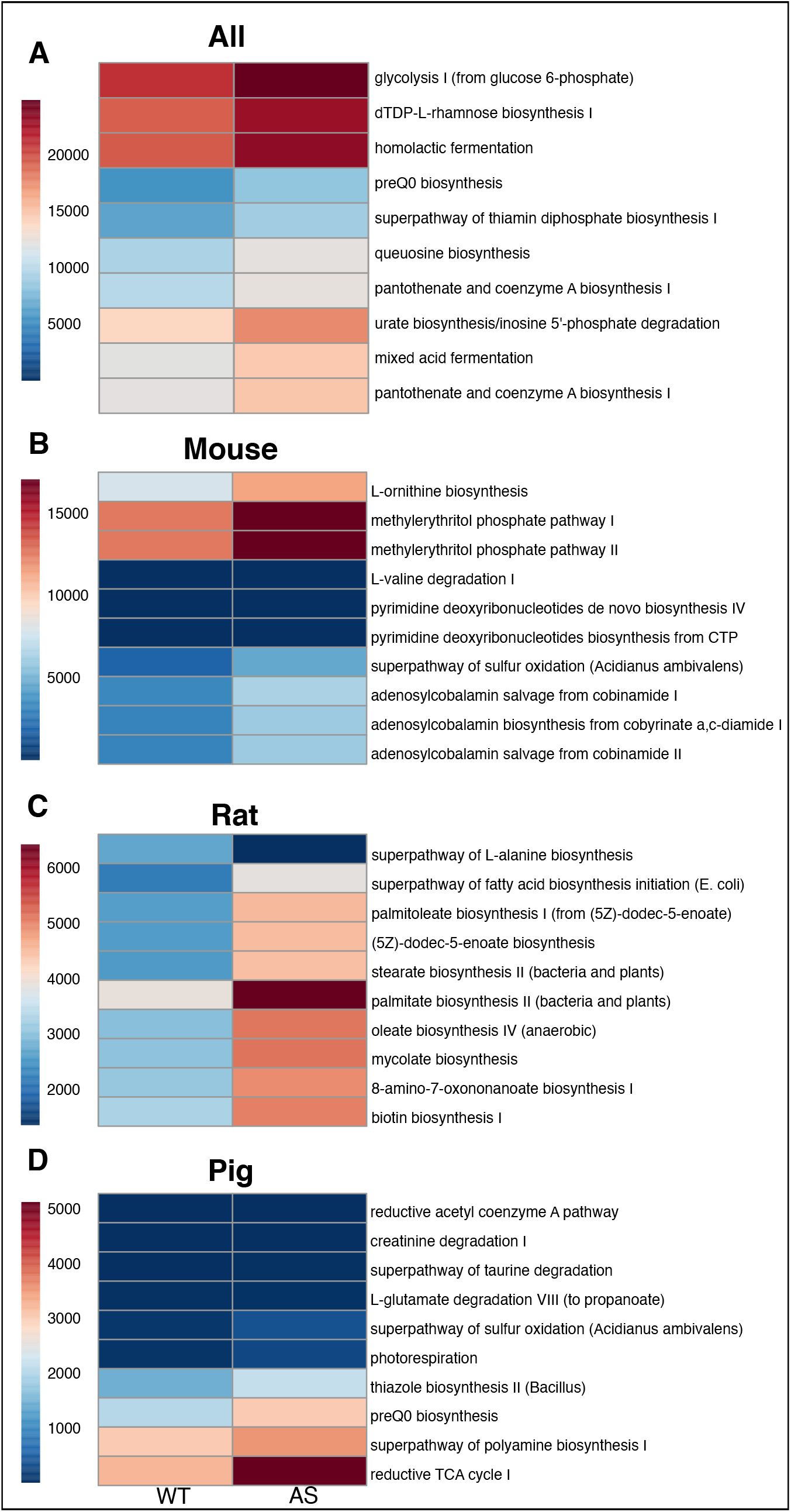
Metabolic pathway prediction analysis shows significant differences between genotype and animal model. Prediction of metabolic pathways based on the microbial ecosystem was analyzed and pathways significantly different in overall (A) and individual animal models (mouse [**A**], rat [**B**], pig [**C**]) showed no overlap but similar functional correlations related to vitamin synthesis and utilization.

In the mouse, metabolic pathways reflect changes based on genotype. Increased activity of amino acid biosynthesis, isopentenyl diphosphate synthesis, adenosylcobalamin salvage, and biosynthesis from vitamin B12 analogs (cobinamides) were found in AS mice compared to WT controls (**Fig 6B**). In the rat, similar increases in the biosynthesis of amino acids, lipokine biosynthesis (palmitoleate), fatty acid synthesis, and biotin biosynthesis were observed (**Fig 6C**). Finally, in the pig model, increase biosynthesis of thiazole (vitamin B1), production of polyamines, and changes in the reductive TCA cycle (responsible for making organic molecules to produce sugars, lipids, amino acids, etc.) were observed in AS compared to WT animals (**Fig 6D**).

Overall, when we compare the individual metabolic differences within the animal models’ similarities among biosynthesis of amino acids pathways are observed. Additionally, they all present different pathways related to the vitamin B complex and its involvement in biological processes, suggesting a common feature of AS across animal models.

## DISCUSSION

In the past decade, there has been a significant increase in research on AS, from basic to applied research, due in part to a large effort to create preclinical animal models for the purpose of identifying targeted treatments and outcome measures for future clinical trials (45). Given that GI symptoms are commonly seen in patients and can significantly impair their quality of life, the identification of factors that can regulate GI physiology is critical in advancing these goals. The gut microbiome is crucial for the establishment of GI physiology and function, and alterations in the colonization of the gut microbiome are prevalent in neurodevelopmental disorders. Characterizing the impacts of *Ube3a* deletion on the gut microbiome using animal models has the powerful advantage to control for variables that are challenging in humans, including environment and diet. As there are currently no gut microbiome studies reported in AS patients, this study represents the first attempt to characterize the microbiome in AS, using animal models, and to identify pathways that could be targeted to improve GI pathophysiology in patients. Here we use 16S ribosomal DNA amplicon sequencing in AS animal models in three species to uncover common colonization features associated with maternal *Ube3a* deletion.

In the current study, we first looked at the biodiversity at different scales, both within each model and common across all three models. The mouse model was found to be much less rich and diverse in comparison to the other two AS models. The lack of diversity and richness, as measured by alpha diversity, has been reported before in laboratory mice compared to wild *M. domesticus*, and is likely in part due to the standard diet to which they are provided (46). However, based on beta diversity, the mouse microbiota seems to be closer to humans than that of rats (47). No major differences in the microbial structure were observed in AS animal models compared to WT controls, which suggests that the overall primary microbial community remained conserved. Given that only bacteria with a greater than 1% overall abundance were assessed, it is possible that the changes could be occurring in low abundance bacterial groups that were not captured in this analysis. Alterations in less abundant bacteria can potentially change interactions within the gut without affecting the overall microbial community.

Findings from the highly abundant taxonomic groups suggest that the main differences between the AS animals and WT controls reside in a reduction of lactic acid bacteria that are essential in the gut for maintaining health (48–51). The decrease of *Bifidobacterium* coupled with an increase in the abundance of *Bacteroides* was seen in AS animals, which has been previously observed in patients with chronic constipation (52, 53). As typically seen in constipation studies, there is no direct consensus whether these changes in the gut microbiome are causal or the result of a side effect of the condition. However, constipation is the most common symptom seen in AS patients, and these findings support a change in the microbial composition associated with the AS genotype. Observing these trends in taxonomic groups within three different animal models of AS further supports that these differences arise due to genetic deletions resulting in AS.

The GI tract communicates bidirectionally with the brain and is closely associated with neurodevelopment, as both develop during early neonatal life in multiple animals. Altered colonization of the gut microbiota, termed “dysbiosis”, has been observed to correlate with disease in patients with neurodevelopmental delays such as autism spectrum disorder (ASD) (13). In ASD, behavioral and neurodevelopmental changes have been correlated with a reduction of *bifidobacterium* and *blautia* species (54), similar to AS animal models seen here, when compared to WT controls. These findings suggest that these microbial community impairments may serve as the main contributor to neurodevelopmental delays. The abundance of other bacterial species, including *desulfovibrio, lactobacillus*, and *bacteroides* are also increased in ASD (54, 55). Our fold change analysis presented an overall increased prevalence of *Lachnospiraceae insertae sedis, Desulfovibrio* and *Odoribacter* in AS in comparison to WT controls. Increased *Lachnospiraceae insertae sedis* has been associated with multiple diseases, including major depressive disorder, and non-alcoholic fatty liver disease (56). Moreover, *Desufovibrio* has been correlated with Parkinson’s disease and its abundance in the gut is directly correlated with disease severity (57). Furthermore, *Odoribacter* has been correlated with attention-deficit/hyperactivity disorder and destabilizes the levels of dopamine and serotonin in the gut (58). This suggests the observed bacterial groups increased in AS align with current studies of other neurodevelopmental diseases (59).

Prediction of the metabolic function changes resulting from the altered gut microbiota in AS animals allow us to understand the metabolic implication of dysbiosis in AS. As explored in the inferred metabolic pathways analysis using PICRUST2, changes in vitamin B12 synthesis and utilization are impacted due to microbial dysbiosis. Since vitamin B12 has the potential to break down homocysteine, increased B12 will increase homocysteine which is directly correlated with dementia, heart disease, and stroke (60). In mice, deficiency of B12 is associated with protection against colitis (61), while in rats, B12 deficiency causes intestinal barrier defects (62). Cobalamin or B12 deficiency can cause increased homocysteine, or hyperhomocysteinemia, which occurs commonly in patients with inflammatory bowel disease (63). Similar changes in the gut microbial ecosystem in ASD studies showed the implications of gut dysbiosis in the production and utilization of vitamins such as B12 (64). The use of comparisons between AS and ASD that lead to microbial dysbiosis and metabolic disparities has the potential to identify what changes in the metabolome and microbiome contribute to disease severity.

In conclusion, the microbial composition analysis of AS within three separate animal models shows a prominent change in the composition and metabolic capacity of the gut microbiome compared to WT control animals. Bacterial groups that are significantly altered within the AS models have also been correlated with other neurodegenerative and GI diseases, highlighting their important role in gut-brain communication. It remains to be determined whether the gut microbiome is a cause or effect of the GI AS symptoms, but the current analysis suggests that the microbial ecosystem may promote adverse gut-brain pathways. Beneficially modulating the gut microbiome may serve to improve both neural and gastric symptomatology in patients with AS, possibly improving overall quality of life.

## ACKNOWLEDGEMENTS

16S rRNA sequencing was carried out by the Host Microbe Systems Biology Core at UC Davis Genome Center.

## FUNDING

- P50HD103526/Eunice Kennedy Shriver National Institute of Child Health and Human Development (PI Abbedutto)
- R01NS097808/NS/NINDS NIH HHS/United States
- Foundation for Angelman Syndrome for Therapeutics (FAST; UB; DJS and JLS)
- UC Davis PREP postbac award (NIHR25GM116690) (BVC)

